# Determinants in the HTLV-1 capsid major homology region that are critical for virus particle assembly

**DOI:** 10.1101/2024.04.07.588439

**Authors:** Huixin Yang, William G. Arndt, Wei Zhang, Louis M. Mansky

## Abstract

The Gag protein of retroviruses is the primary driver of virus particle assembly. Particle morphologies among retroviral genera are distinct, with intriguing differences observed relative to HIV-1, particularly that of human T-cell leukemia virus type 1 (HTLV-1). In contrast to HIV-1 and other retroviruses where the capsid (CA) carboxy-terminal domain (CTD) possesses the key amino acid determinants involved in driving Gag-Gag interactions, we have previously demonstrated that the amino-terminal domain (NTD) encodes the key residues crucial for Gag multimerization and immature particle production. Here in this study, we sought to thoroughly interrogate the conserved HTLV-1 major homology region (MHR) of the CA_CTD_ to determine whether this region harbors residues important for particle assembly. In particular, site-directed mutagenesis of the HTLV-1 MHR was conducted, and mutants were analyzed for their ability to impact Gag subcellular distribution, particle production and morphology, as well as the CA-CA assembly kinetics. Several key residues (i.e., Q138, E142, Y144, F147 and R150), were found to significantly impact Gag multimerization and particle assembly. Taken together, these observations imply that while the HTLV-1 CA_NTD_ acts as the major region involved in CA-CA interactions, residues in the MHR can impact Gag multimerization, particle assembly and morphology, and likely play an important role in the conformation the CA_CTD_ that is required for CA-CA interactions.

## Introduction

Human T-cell leukemia virus type 1 (HTLV-1) is in the deltaretrovirus genus of the *Retroviridae*. HTLV-1 was the first infectious, oncogenic human retrovirus discovered [1, 2]. Other HTLV subtypes have been subsequently identified over the years, including HTLV-2 [3], HTLV-3 [4-6], and HTLV-4 [6]. HTLV-2 was first identified in a patient with hairy cell leukemia [3], while HTLV-3 and HTLV-4 were discovered in bushmeat hunters [6]. HTLV-1 is clinically associated with adult T-cell leukemia/lymphoma (ATLL) and HTLV-1-associated myelopathy/tropical spastic paraparesis (HAM/TSP) [7-9]. HTLV-1 infection has a high prevalence in particular regions of the world, including Japan, the Caribbean, Central and South America, Africa, and the Middle East [10]. To date, it is estimated that 15 to 20 million individuals are infected with HTLV-1 worldwide [11, 12], and approximately 10% of those infected individuals develop ATL or TSP/HAM [13].

The development of antiretroviral drugs directed against different steps in the viral life cycle of human immunodeficiency virus type 1 (HIV-1) have saved millions of lives and led to a remarkable reduction in overall morbidity and mortality when used in combination therapy [14]. Reverse transcriptase, protease and the integrase enzymes represent the major antiretroviral targets, along with inhibitors directed against virus fusion. Any step in the virus life cycle could be a potential antiretroviral target, providing the basis for next generation antiretroviral development. For example, virus particle assembly is a critical step that has been exploited in the development of a capsid assembly inhibitor, lenacapavir [15-17]. Lenacapavir was approved as a long-acting antiretroviral drug. In contrast to these achievements with HIV-1, the development of treatments and therapies directed against HTLV-1 has lagged behind, which is due in part to a lack of detailed information the HTLV-1 life cycle [18, 19]. Therefore, there remains a strong need for basic research investigation of HTLV-1 replication in order to identify promising antiviral targets for therapeutic intervention as well as for vaccine development.

The HTLV-1 Gag polyprotein is a 53 kDa protein that contains three subdomains: matrix (MA), capsid (CA) and nucleocapsid (NC) [20]. The CA domain encodes the majority of the critical amino residues required for Gag multimerization that drive Gag-Gag interactions and ultimately particle assembly. The CA domain is divided into two structurally distinct subdomains, the capsid amino-terminal domain (CA_NTD_) and the capsid carboxy-terminal domain (CA_CTD_), which are connected by a flexible inter-domain region. We have previously demonstrated that HTLV-1 CA_NTD_ encodes the key residues (*i.e.*, M17 and Y61) crucial for Gag multimerization and immature particle production [21]. However, the specific role of HTLV-1 CA_CTD_ in particle assembly was not clearly defined. Within the CA_CTD_ resides the highly conserved major homology region (MHR), which is comprised of 20 amino acid residues. The MHR region has been extensively studied among other different retroviruses, including HIV-1 [22-31], Rous sarcoma virus (RSV) [31-34], Mason-Pfizer monkey virus (MPMV) [35], and murine leukemia virus (MLV) [36]. These studies identified specific residues in the MHR that play a crucial role in the assembly of the immature Gag lattice for various retroviruses (*e.g.*, for HIV-1 CA, Q155, K158, Y164 and R167) [22, 25-28, 30]. However, to date, no systematic analysis has been conducted for the HTLV-1 CA MHR. Previously, the only residue in the HTLV-1 MHR region that was reported to be critical for virus assembly was E142 [37].

In this study, we sought to thoroughly interrogate the conserved HTLV-1 MHR region of the CA_CTD_ in order to determine whether this region harbors residues important for particle assembly. To achieve this goal, we conducted alanine-scanning mutagenesis of the HTLV-1 CA MHR region. In particular, a panel of 20 site-directed mutants were created and were analyzed for their ability to impact Gag subcellular distribution, particle production and morphology, as well as the CA-CA assembly kinetics. Several key residues (*i.e.*, Q138A, E142A, Y144A, F147A, and R150A) were found to significantly impact Gag multimerization as well as particle assembly. Taken together, these observations provide the first detailed insights into the critical role of the HTLV-1 CA MHR and imply that residues in HTLV-1 CA MHR can impact Gag multimerization, particle assembly and morphology, and likely play an important role in the conformation of the CA_CTD_ that is required for the HTLV-1 CA_NTD_ to drive CA-CA interactions.

## Results

### Identification of HTLV-1 CA MHR residues that impact particle production

Given previous observations that residues in retroviral CA MHR region play a critical role in viral particle assembly, including HIV-1 [22-31], RSV [31-34], MPMV [35] and MLV [36], we sought to address this knowledge gap with HTLV-1 CA by conducting alanine-scanning mutagenesis of the entire HTLV-1 CA MHR domain (**Figure 1**). In order to analyze particle production, we utilized a previously characterized tractable HTLV-1-like particle system to assess the impact of individual mutations on authentic, immature virus particle assembly and release [20, 38, 39]. With the exception of E142 [37], 19 of the 20 amino acid residues in the MHR domain have not been previously characterized for HTLV-1.

**Figure 1.**
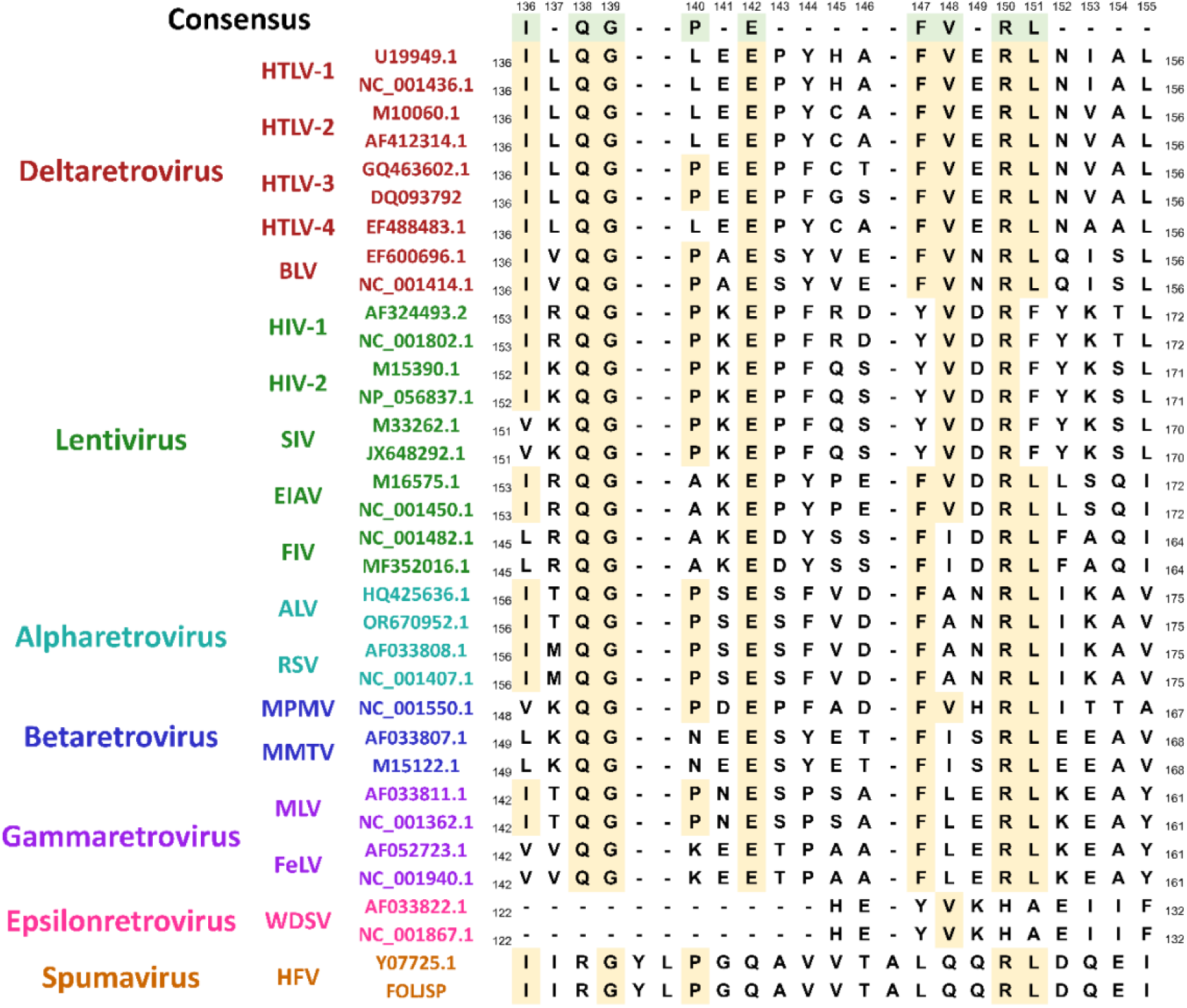
Amino acid alignment of the major homology region in the capsid carboxy-terminal domain of Gag among various retroviral genera. The major homology region (MHR) in the capsid carboxy-terminal domain (CA_CTD_) from the Gag polyprotein of 34 different retroviruses (in 7 different genera) were aligned and analyzed by using SnapGene (GSL Biotech, San Diego, CA). Isolates from different retroviral genera are indicated. The retrovirus MHR sequences analyzed included human T-cell leukemia virus type 1 (HTLV-1; GenBank accession number U19949.1 and NC_001436.1), human T-cell leukemia virus type 2 (HTLV-2; GenBank accession number M10060.1 and AF412314.1), human T-cell leukemia virus type 3 (HTLV-3; GenBank accession number GQ463602.1 and DQ093792.1), human T-cell leukemia virus type 4 (HTLV-4; GenBank accession number EF488483.1), bovine leukemia virus (BLV; GenBank accession number EF600696.1 and NC_001414.1), human immunodeficiency virus type 1 (HIV-1; GenBank accession number AF324493.2 and NC_001802.1), human immunodeficiency virus type 2 (HIV-2; GenBank accession number M15390.1 and NP_056837.1), simian immunodeficiency virus (SIV; GenBank accession number M33262.1 and JX648292.1), equine infectious anemia virus (EIAV; GenBank accession number M16575.1 and NC_001450.1), feline immunodeficiency virus (FIV; GenBank accession number NC_001482.1 and MF352016.1), avian leukosis virus (ALV; GenBank accession number HQ425636.1 and OR670952.1), Rous sarcoma virus (RSV; GenBank accession number AF033808.1 and NC_001407.1), Mason-Pfizer monkey virus (MPMV; GenBank accession number NC_001550.1), mouse mammary tumor virus (MMTV; GenBank accession number AF033807.1 and M15122.1), murine leukemia virus (MLV; GenBank accession number AF033811.1 and NC_001362.1), feline leukemia virus (FeLV; GenBank accession number AF052723.1 and NC_001940.1), walleye dermal sarcoma virus (WDSV; GenBank accession number AF033822.1 and NC_001867.1), and human foamy virus (HFV; GenBank accession number Y07725.1 and FOLJSP). The HTLV-1 CA numbers are indicated at the top of the figure. The amino acid numbering to the left and right of each MHR sequence corresponds to the CA protein numbering from the respective retrovirus. The consensus sequence with a threshold of >50% is indicated at the top (amino acids highlighted in green), while the amino acids that match the consensus among the various MHR sequences are indicated by yellow highlighting.

Alanine-scanning mutants of the 20 amino acid residues in the HTLV-1 CA MHR were analyzed for their impact on particle production by immunoblot analysis, and five of these mutants (*i.e.*, Q138A, E142A, Y144A, F147A and R150A) were found to result in a 2-fold or greater reduction in particle production relative to that of WT (**Figure 2**). In particular, the HTLV-1 CA Q138A and Y144A mutants reduced particle production by 5.4-fold and 3.2-fold, respectively, compared to WT (**Figure 2A**), and the CA F147A and R150A mutants reduced particle production by 2.4-fold and 2.9-fold, respectively, compared to WT (**Figure 2A**). While comparable results have not been previously reported for Q138 and Y144 in regards to particle production, observations made with F147A and R150A are consistent with comparable mutations in HIV-1 CA (*i.e.,* Y164A and R167A, respectively) in the HIV-1 CA MHR domain on particle production, in which both mutants were found to result in particle reductions [22, 26, 30]. The HTLV-1 CA E142A mutant reduced particle production by 2.6-fold compared to WT (**Figure 2A**), which is consistent with the literature, where HTLV-1 CA E142K mutant led to a comparable 2.3-fold decrease in particle production [37].

**Figure 2.**
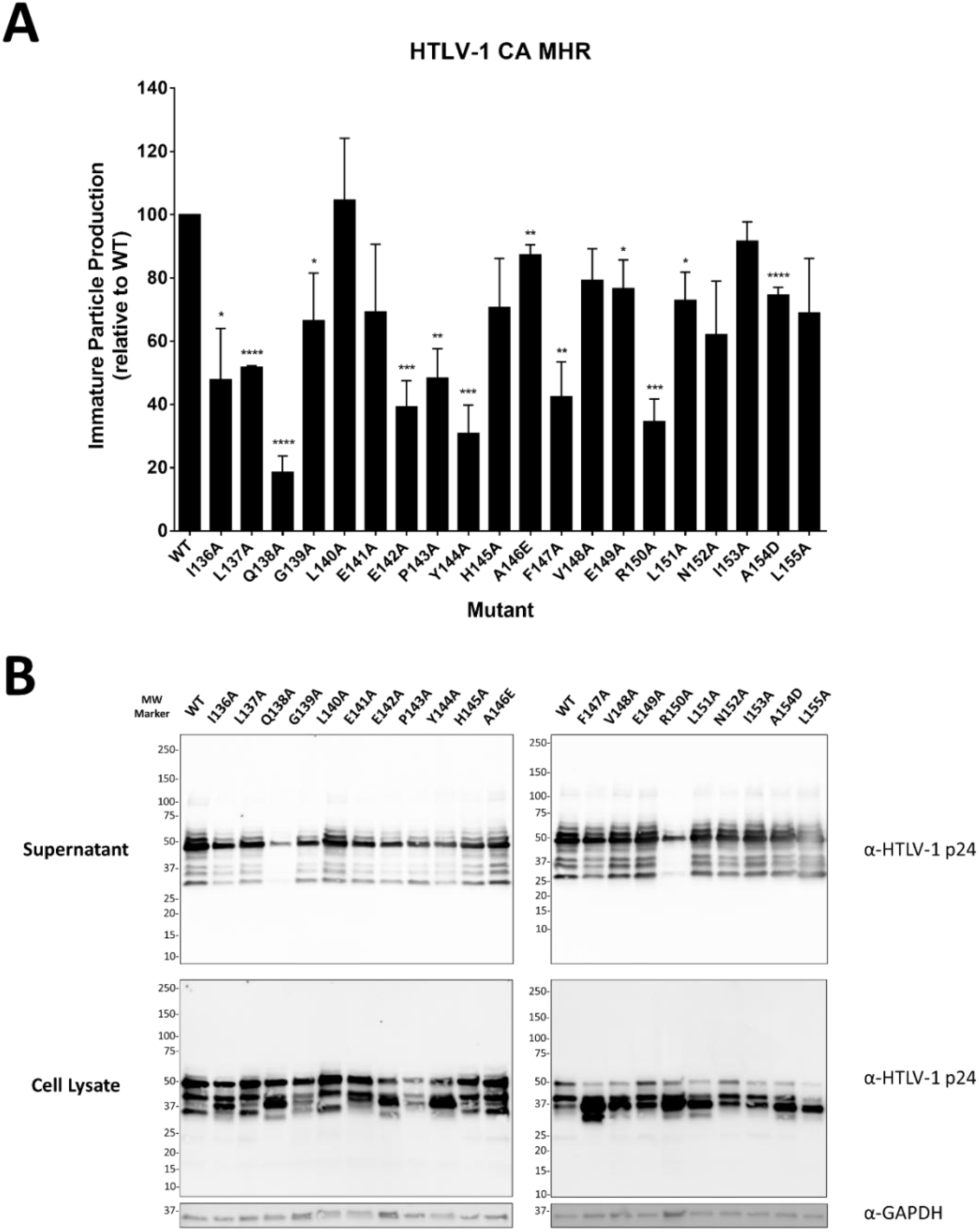
Particle production of HTLV-1 capsid protein major homology region mutants. Site-directed mutagenesis of amino acid residues located in the major homology region (MHR) of the capsid carboxy-terminal domain (CA_CTD_) were created and the mutants were analyzed for their ability to produce particles relative to that of the WT. Transfection of 293T/17 cells with plasmids expressing the HTLV-1 Gag (WT or alanine-scanning mutant) was done and 48 h post-transfection cell culture supernatants and cell lysates were harvested. (A) Analysis of particle production. Immunoblot analysis was conducted to determine particle production of mutants relative to that of WT HTLV-1 Gag (set to 100). Data was collected from three independent experiments. Error bars represent the standard error of the mean. Significance relative to WT was determined by using an unpaired t-test. ****, *P* < 0.0001; ***, *P* < 0.001; **, *P* < 0.01; *, *P* < 0.05. (B) Immunoblot analysis. Shown is a representative immunoblot analysis by using an anti-HTLV CA antibody of the Gag protein detected from both cell culture supernatant and from cell lysates. Detection of GAPDH from the cell lysates was used to normalize detection of WT Gag and that of the MHR mutants.

### HTLV-1 CA MHR mutants alter HTLV-1 particle morphology

Critical residues in the HTLV-1 CA_NTD_ (*i.e.*, M17 and Y61) have been reported to impact particle morphology, implicating that the HTLV-1 CA_NTD_ can cause defects in Gag to multimerization, lattice structure, and infectious particle formation [21]. However, a current knowledge gaps exists regarding the role of HTLV-1 CA_CTD_ on particle morphology. Therefore, we sought to investigate the impact of HTLV-1 CA MHR mutants on immature particle morphology. Selected HTLV-1 CA MHR mutants (*i.e.,* Q138A, E142A, Y144A, F147A, and R150A) were examined for immature particle morphology compared to that of parental HTLV-1 WT immature particles (**Figure 3**).

**Figure 3.**
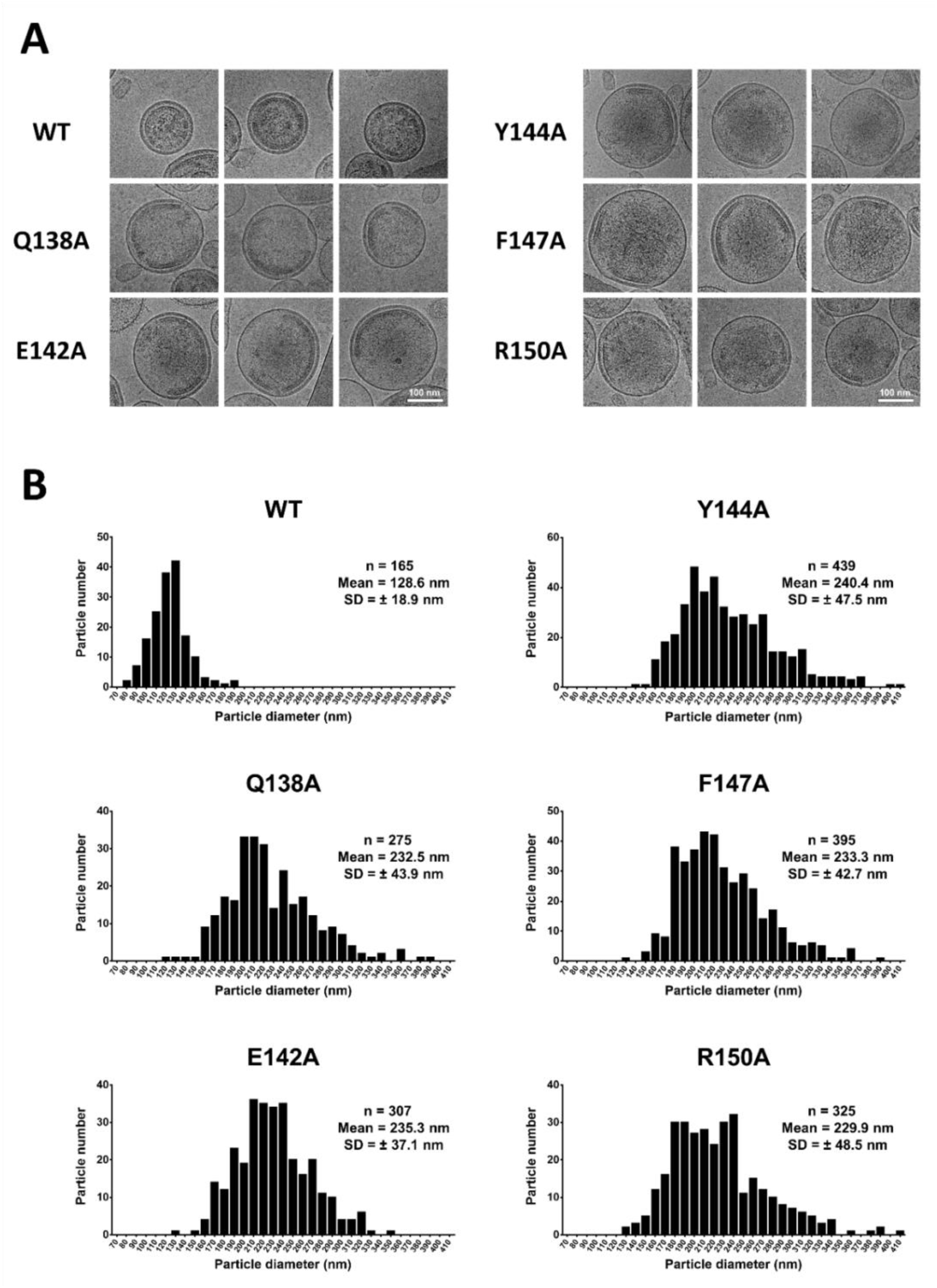
Cryo-electron microscopy of immature HTLV-1 particles with mutation of the capsid protein major homology region. Selected HTLV-1 Gag mutants in the capsid (CA) major homology region (MHR) (*i.e.*, Q138A, E142A, Y144A, F147A, and R150A) were transiently transfected into 293T/17 cells, and the particles produced from the cells were collected from cell culture supernatants, and concentrated and purified by ultracentrifugation prior to cryo-electron microscopy (cryo-EM) analysis. (A) Cryo-EM analysis. Representative images are shown for WT and the CA MHR mutants (scale bar, 100 nm). (B) Size distributions of WT and CA MHR mutant particles. Particle diameters were determined by averaging the measurements in two perpendicular directions. The number (n) of particles, mean, and standard deviation (SD) are indicated.

Cryo-EM analysis of WT HTLV-1 immature particles revealed regions of flat Gag lattice electron density that did not fully follow the curvature of the spherical lipid bilayer (**Figure 3A**), which is consistent to previous reports [21]. HTLV-1 WT immature particles were primarily spherical, with some particles having different morphologies. In contrast, HTLV-1 CA MHR mutants were found to have an increase in non-spherical morphologies with clear gaps and defects in the immature Gag lattice. Furthermore, the HTLV-1 CA MHR mutants were more commonly observed to form flat lattices compared to that of WT. The quantification of WT HTLV-1 particles revealed an average diameter of 128.6 nm (SD = 18.9 nm) (**Figure 3B**). The mean particle diameter was significantly larger for the HTLV-1 CA MHR mutants (*i.e.*, Q138A, E142A, Y144A, F147A, and R150A) than that of the HTLV-1 WT immature particles, with an average diameter of 232.5 nm (SD = 43.9 nm), 235.3 nm (SD = 37.1 nm), 240.4 nm (SD = 47.5 nm), 233.3 nm (SD = 42.7 nm) and 229.9 nm (SD = 48.5 nm), respectively, (**Figure 3B**). Taken together, these observations imply that the HTLV-1 CA MHR mutants impose a defect in Gag multimerization to form a proper Gag lattice structure.

### Perturbation of Gag subcellular distribution

Gag subcellular distribution is an efficient method to qualitatively assess the ability of the mutant Gag proteins to multimerize to form Gag puncta. To determine if the subcellular distribution of HTLV-1 mutants was associated with reductions in particle production and particle morphology, the selected MHR mutants (*i.e.*, Q138A, E142A, Y144A, F147A, and R150A) were engineered an in-frame YFP tag at the carboxy-terminus of the Gag protein (**Figure 4**). The parental HTLV-1 WT and mutant Gag-YFP constructs were transfected into HeLa cells (**Figure 4A**) and analyzed for their ability to multimerize and form Gag puncta at 24h posttransfection (**Figure 4B**), where puncta formation at the plasma membrane was indicative of assemblies that may result in particle production. The lack of Gag puncta and the presence of diffuse Gag-eYFP fluorescence in transfected cells was interpreted as an indication of low or no Gag multimerization, which would negatively impact particle release.

**Figure 4.**
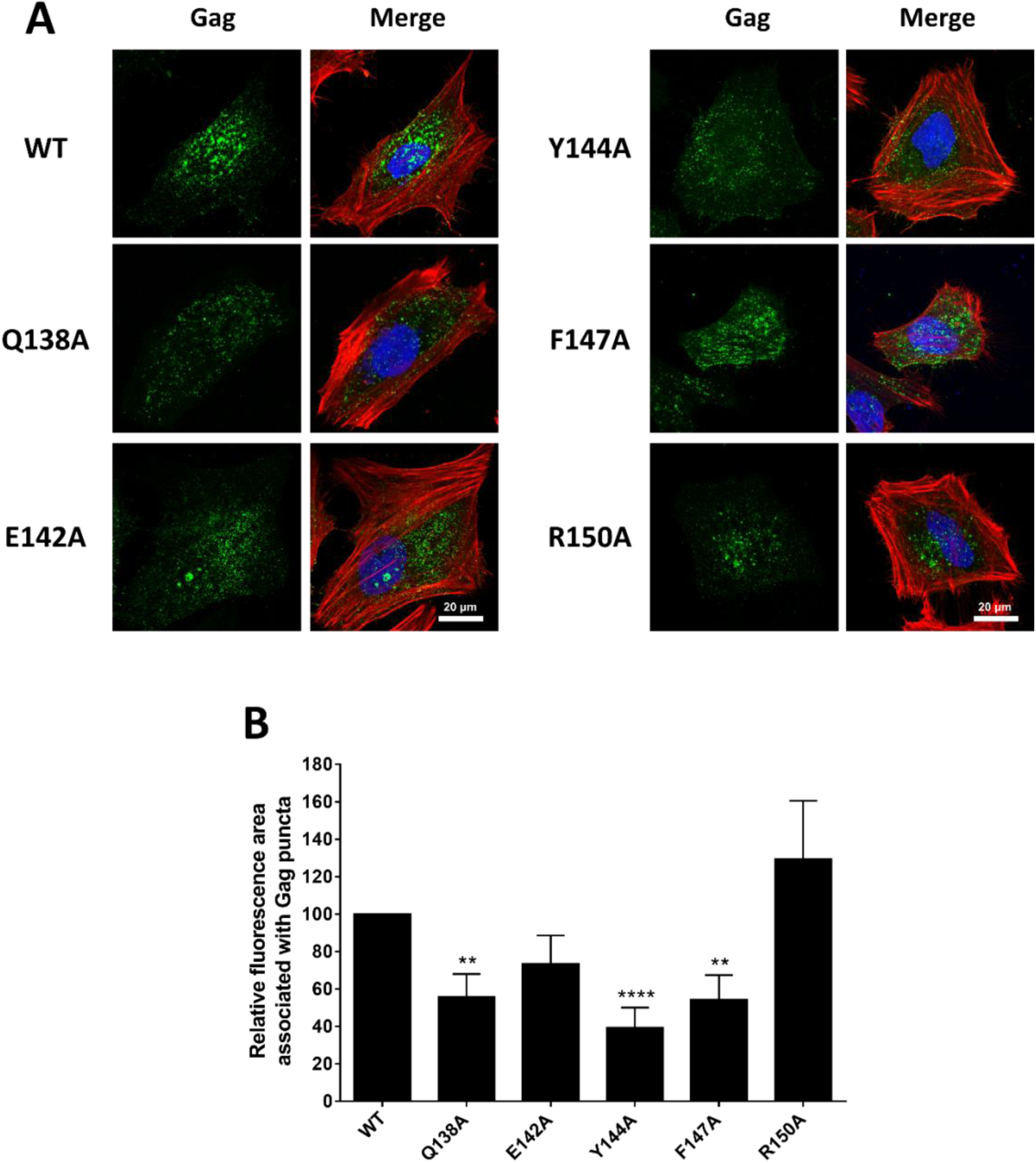
Subcellular localization of the HTLV-1 Gag mutants with mutation of the capsid protein major homology region. Selected HTLV-1 Gag mutants in the capsid (CA) major homology region (MHR) (*i.e.*, Q138A, E142A, Y144A, F147A, and R150A) were transiently transfected into HeLa cells and imaged 24 h post-transfection. (A) Subcellular localization of HTLV-1 Gag mutants. Representative confocal images of HTLV-1 CA MHR mutants are shown. At least 15 individual cells were imaged across three independent replicates for a total of 15 cells. Scale bar, 20 μm. Gag (green, eYFP fluorescence), nuclei (blue; stained with DAPI), and actin (red, stained with actin red) are indicated. (B) Analysis of HTLV-1 Gag puncta formation. The mean relative fluorescence area associated with Gag puncta, relative to that of WT, was determined as described in Materials and Methods. Data was collected from three independent experiments. Error bars represent the standard error of the mean. Significance relative to that of WT was determined by using an unpaired t-test. ****, *P* < 0.0001; **, *P* < 0.01.

The efficiency of HTLV-1 Gag puncta formation was analyzed as described in Materials and Methods. The E142A and R150A mutants were observed to result in Gag puncta formation at levels comparable to that of WT, with a mean of 73% and 129%, respectively (**Figure 4B**). However, the Q138A, Y144A and F147A mutants resulted in significantly reduction in Gag puncta formation (with a mean of 56%, 39% and 54%, respectively) compared to WT Gag (**Figure 4B**), indicating that the Q138A, Y144A and F147A mutations led to reduced Gag puncta formation. Taken together, these observations indicate that mutation of the HTLV-1 CA MHR impacts HTLV-1 Gag subcellular distribution.

### Disruption of HTLV-1 CA in vitro assembly

*In vitro* protein assembly assays have been very useful for analyzing CA protein assembly due to this being an efficient system to perturb protein-protein interactions in the absence of cellular effects. To assess the impact of selected HTLV-1 CA MHR mutations on protein interactions, a HTLV-1 CA construct was generated to express and purify recombinant protein for *in vitro* assembly assays (**Figure 5**). HTLV-1 CA WT and selected MHR mutants (*i.e.*, Q138A, E142A, Y144A, F147A and R150A, based on the amino acid numbering in the Gag protein) were dialyzed against assembly buffer to induce helical assembly formation and quantified by turbidity measurements at A_340_ (**Figure 5A-5F**). At this wavelength, the turbidity measurement reflects the state of protein assembly from a soluble protein state to higher-order CA assemblies. The ability of the Q138A, E142A, Y144A, F147A and R150A mutants to form helical assemblies was measured by taking readings every 5 min over a 1 h time frame.

**Figure 5.**
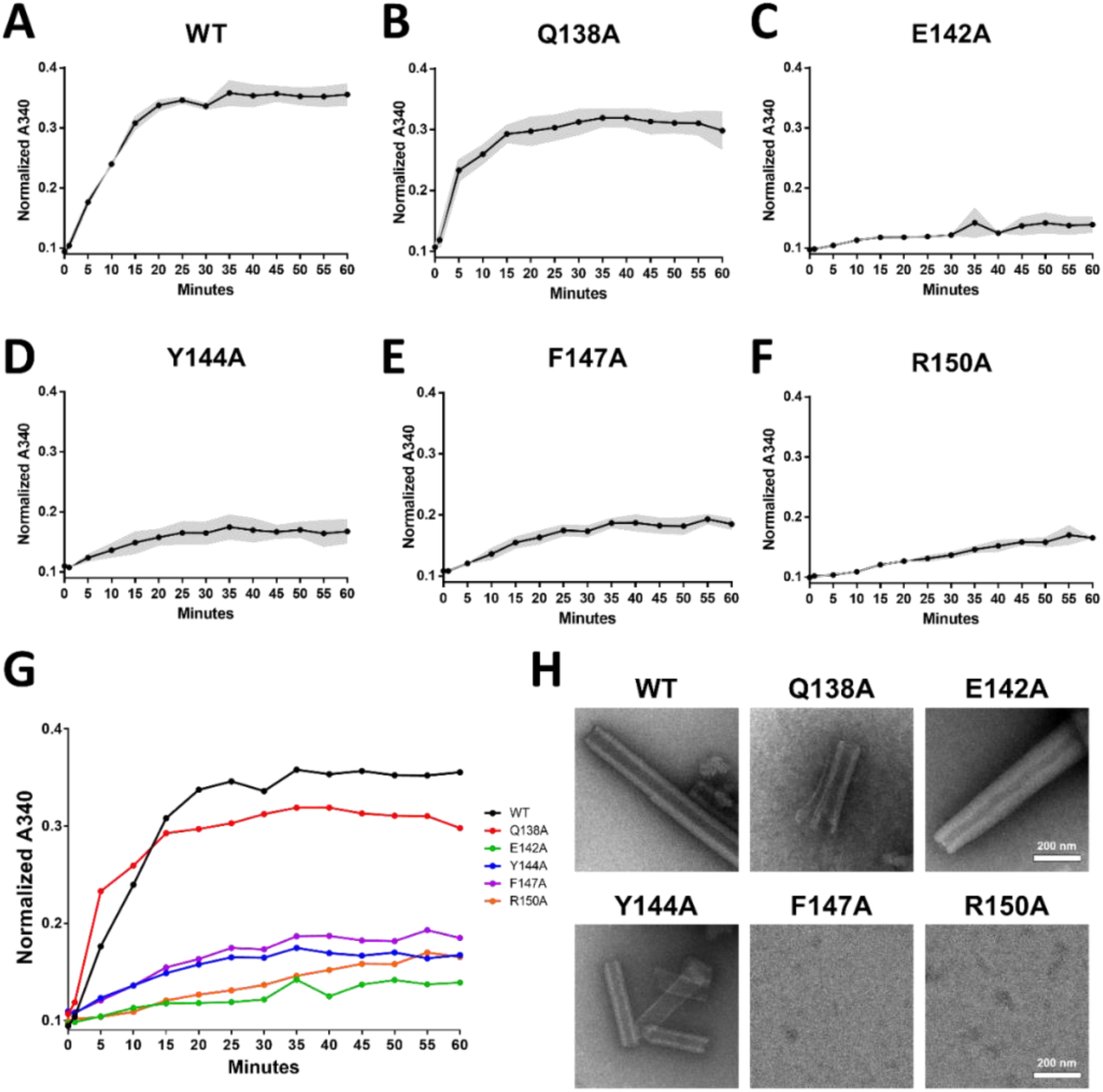
Cell-free assembly of HTLV-1 capsid protein tubular assemblies. (A-F) Assembly kinetics of HTLV-1 capsid (CA) major homology region (MHR) mutants (*i.e.*, Q138A, E142A, Y144A, F147A, and R150A) were monitored by measuring light scattering at 340 nm. The CA MHR mutant assemblies were pre-dialyzed into assembly buffer and assembly was initiated by the addition of IP6. Readings were taken every 5 min for 60 min and corrected for any changes in reaction volume due to dialysis. The grey area shown in each panel represents the average ± standard deviation from three independent experiments. (G) CA MHR mutant assembly kinetics. Shown is the analysis of the CA MHR mutant assembly kinetics compared to WT (H) Representative electron microscopy images of negatively stained HTLV-1 CA MHR mutant assembly products, which were sampled at 60 min. Scale bar, 200 nm.

The normalized A_340_ reading for HTLV-1 WT (**Figure 5A**) peaked at 25 min (*i.e.*, A_340_ = 0.35), and then remained stable. HTLV-1 Q138A (**Figure 5B**) had similar trends with WT, but with lower A340 nm values (*i.e.*, A_340_ = 0.30) compared to WT. The trend of normalized A_340_ values of HTLV-1 E142A, Y144A, F147A, and R150A, were different from WT and Q138A, which were relatively stable and slightly increased, but all had low turbidity readings (*i.e.*, A_340_ = 0.14, 0.17, 0.19 and 0.17, respectively) (**Figure 5C-5F**) during the 1 h time course. The Q138A mutant had a slight reduction in assembly kinetics relative to that of WT, while the E142A, Y144A, F147A and R150A CA mutants had severe reductions in assembly kinetics relative to that of WT. After completion of the 1h incubation, the Q138A, E142A, Y144A, F147A and R150A CA mutants led to significant reductions in turbidity relative to that of WT, which were 1.2-fold, 2.6-fold, 2.1-fold, 1.9-fold and 2.1-fold lower than that of WT, respectively (**Figure 5G**). Taken together, these observations indicated that the selected HTLV-1 MHR mutants analyzed led to a disruption of HTLV-1 CA assembly in the *in vitro* assay and provides evidence that amino acid residues in the MHR play critical roles in CA-CA interactions.

The visualization of HTLV-1 CA helical assemblies by negative stain TEM revealed that HTLV-1 WT CA assembled into stable, regularly ordered tubular assemblies, while the Q138A, E142A and Y144A mutants were capable of forming stable, regularly ordered tubular assemblies, but with lower efficiency (**Figure 5H**). The HTLV-1 F147A and R150A MHR mutants resulted in protein aggregations without forming regularly ordered uniform tubular assemblies (**Figure 5H**). Taken together, these observations support the conclusion that the HTLV-1 MHR mutants, particularly F147A and R150A, can cause defects in protein-protein interactions that prevent helical tube assembly formation.

## Discussion

While a number of studies have been conducted on the MHR, there remains many unanswered questions regarding the evolutionary conservation and function of the MHR [26, 40]. Models have been previously proposed to help explain MHR conservation and function. One model is that the MHR region plays an essential role in forming a stable and conserved structure for the retroviral CA monomer. In particular, Zurowska et al. proposed that the HIV-1 MHR structure, characterized by a densely packed strand-turn-helix configuration, could help to elucidate its function [41]. Within the HIV-1 CA_CTD_, this configuration is stabilized by a network of hydrogen bonds and salt bridges between the side chains of glutamine, glutamate, and arginine. This network appears to be absolutely conserved within the retroviral MHR motif [40-42]. Moreover, conserved positions are occupied by large hydrophobic side chains — typically phenylalanine, tyrosine, or leucine — which constitute part of the hydrophobic core of CA_CTD_. However, distinctive differences exist among retroviruses. For example, the HIV-1 MHR residue K158 plays a role in stabilizing the six-helix bundle [43-46]. In contrast, no residues in the HTLV-1 MHR have been clearly demonstrated to serve this function. The MHR region may also possess additional functions. Tanaka et al. concluded that the conserved residues function after multimerized HIV-1 Gag has associated with genomic RNA, recruited critical host factors involved in assembly, and has been targeted to the plasma membrane (PM) [26]. Koma et al. have previously suggested that MHR mutations can significantly impact the ability of Gag to multimerize at the PM [30].

The HTLV-1 CA_NTD_ has been previously shown to encode key residues crucial for Gag multimerization and immature particle production [21]. However, there is limited information available on the role of HTLV-1 CA_CTD_, specifically the MHR region, in particle assembly, morphology, and its contribution to CA-CA interactions. In this study, we sought to thoroughly interrogate the conserved HTLV-1 MHR of the CA_CTD_ in order to determine whether this region harbors residues important for particle assembly. To address this knowledge gap, we conducted a panel of twenty site-directed mutant spanning all HTLV-1 MHR residues and analyzed their impact on particle production, particle morphology, Gag subcellular distribution, and on formation of CA assemblies *in vitro*. To the best of our knowledge, this is the first comprehensive comparative study of the HTLV-1 MHR region. Previously, only the E142 residue has been analyzed for its influence of virus particle assembly [37].

Our alanine-scanning mutagenesis of the MHR revealed five key residues (Q138, E142, Y144, F147, and R150) that had a significant impact HTLV-1 particle production (**Figure 2**) and morphology (**Figure 3**). Furthermore, several mutants were identified that reduced Gag puncta formation (**Figure 4**) as well as the formation of CA assemblies *in vitro* (**Figure 5**). With the exception of Y144, the other 4 amino acid residues are conserved among retroviruses. Together, these observations demonstrate that several of the conserved residues are important in HTLV-1 assembly.

The HTLV-1 Q138 is an important residue in the HTLV-1 MHR since Q138A significantly reduced particle production (**Figure 2**), altered virus particle morphology (**Figure 3**), and decreased the efficiency of Gag puncta formation (**Figure 4**). Similar effects have been observed with the corresponding amino acid residue in the HIV-1 MHR (*i.e.,* Q155) [23-25] and in the RSV MHR (*i.e.,* Q158) [32, 33]. Mutagenesis of the RSV Q158 residue revealed that substitution to a variety of other amino acid residues (*i.e.,* Q158E, Q158L, Q158N, Q158P, and Q158R) led to clear defects in particle production [32, 33]. Intriguingly, we observed that the HTLV-1 Q138A forms stable, uniformly ordered tubular CA assemblies with slightly lower efficiency than WT (**Figure 5**). This is in contrast to HIV-1 (Q155A), where the comparable substitution results in very low levels of CA assemblies *in vitro* and blocks HIV-1 particle assembly [23-25].

In addition to HTLV-1 Q138, the HTLV-1 E142 residue was found to be an important conserved residue in the HTLV-1 MHR, as the E142A mutant resulted in a 2.6-fold reduction in particle production (**Figure 2**) and altered particle morphology (**Figure 3**). Our observations are similar to that reported by Rayne et al. (2001), in which the HTLV-1 E142K had 2.3-fold reduction in immature particle production than that of WT [37]. We further observed that HTLV-1 E142A mutant led to a disruption in the formation of CA tubular assemblies *in vitro* (**Figure 5**). This is similar that observed for the HIV-1 E159A mutant, which impairs particle assembly and CA assembly *in vitro* [25].

Intriguingly, the HTLV-1 Y144, which is a non-conserved residue among retroviruses disrupts particle production, morphology, and Gag puncta formation (**Figures 2-4**). However, varying observations have been made in creating CA tubular assemblies for HTLV-1 Y144A compared to the comparable residue in the HIV-1 CA (*i.e.,* F161A). The HIV-1 F161A mutation prevented the formation of tubular assemblies [23]. In contrast, we observed that HTLV-1 Y144A led to the formation of stable, regularly ordered and uniform tubular assemblies (**Figure 5H**).

The HTLV-1 F147A mutant in the MHR was also found to reduce particle production, alter particle morphology, reduce puncta formation, and disrupt CA tubular assemblies (**Figure 2-5**). Similar results have been seen with HIV-1, where a comparable mutation (*i.e.,* HIV-1 Y164A) led to severe reductions in virus particle production, altered virion morphology, and reduced Gag multimerization [22, 23, 26, 30, 47]. Intriguingly, the analogous residue (*i.e.,* F167) in HIV-1 and RSV is thought to impact the CA_CTD_-CA_CTD_ interface [48]. Mutation of HTLV-1 R150 to alanine (*i.e.,* R150A) leads to an approximate 3-fold reduction in particle production (**Figure 2**), alters particle morphology (**Figure 3**), disrupts *in vitro* CA protein assembly (**Figure 5F**), and prevents uniform tubular CA protein formation (**Figure 5H**). These observations are comparable to the observations made with the HIV-1 R167A mutant [22, 25], which is critical in determining the conformational stability of the monomeric form of HIV-1 CA_CTD_ [24].

It is notable that not all conserved residues in the HTLV-1 MHR region led to significant reductions in particle production. In particular, mutation of V148 and L151 residues did not impact particle production (**Figure 2**). In our mutagenesis of the HTLV-1 MHR, some phenotypes that were distinct from HIV-1 were observed [23, 29]. For example, the HTLV-1 V148A mutant resulted in modest reduction particle production compared to that of WT (**Figure 2**), while the comparable HIV-1 CA MHR mutation (*i.e.,* V165A) destabilizes HIV-1 CA_CTD_ monomer folding and stability [23]. Additionally, HTLV-1 L151A also resulted in modest reduction in particle production compared to WT (**Figure 2**), while the comparable mutation in HIV-1 (*i.e.,* F168A) significantly reduced particle production [29]. Intriguingly, some residues implicated as being critical for HIV-1 were not important for HTLV-1. For example, HIV-1 K158 residue is vital as an IP6 binding site [44], while the HTLV-1 CA mutation (*i.e.,* E141A) did not significantly impact on immature particle production (**Figure 2**), indicating some distinct differences between HTLV-1 and HIV-1 CA, at least in regards to residues involved in IP6 binding.

Analysis of HTLV-1 MHR mutants clearly identified conserved residues that resulted in phenotypes comparable to that observed with other retroviruses such as HIV-1 and RSV, thereby implicating a conservation of structure and function for selected MHR residues. However, in the present study we identified residues with distinct effects on particle assembly, which differences in structure or functional interactions compared to that of other retroviruses. Structural information of the HTLV-1 CA should help in gaining deeper insights into distinct differences observed with the HTLV-1 MHR (PDB: 1QRJ) [49]. Since exhaustive site-directed mutagenesis of every amino acid residue of the CA_CTD_ has not been conducted to date, in is possible that residues in the intra-hexamer and inter-hexamer interface play a significant role in CA-CA interactions – and therefore should be explored in future research. Analysis of other residues, such as the F147 and R150 residues would be important to determine their role in CA-CA interactions, particle assembly and morphology.

In summary, we have identified several residues (*i.e.,* Q138A, E142A, Y144A, F147A, and R150A) within the HTLV-1 CA_CTD_ MHR that have a major impact Gag multimerization and particle assembly. This highlights the particularly critical impact of residues in the HTLV-1 CA_CTD_ MHR on virus assembly. These findings provide valuable insights into the critical role that HTLV-1 CA MHR plays in the interactions between CA molecules, likely in a structural role that allows for CA-CA interactions. In particular, these observations provide evidence supporting the conclusion that residues in HTLV-1 CA MHR can impact particle assembly and morphology, and likely play an important role in the conformation of the CA_CTD_ that is required for the HTLV-1 CA_NTD_ to drive CA-CA interactions.

## Materials and Methods

### Plasmids, cell lines, and reagents

The HTLV-1 Gag expression plasmids, pN3-Gag and pEYFP-N3-Gag, as well as the HTLV-1 Env expression plasmid, have been previously described [21]. HeLa and HEK293T/17 cells lines were purchased from ATCC (Manassas, VA) and cultured in Dulbecco’s modified Eagle medium (DMEM) supplemented with 10% Cytiva HyClone Fetal Clone III (FC3; Cytiva Life Sciences, Marlborough, MA) and 1% Penicillin-Streptomycin (Pen Strep; Invitrogen, Carlsbad, CA) at 37 °C in 5% CO_2_. All cells used in this study were certified as being free of mycoplasma contamination.

### Site-directed mutagenesis of Gag expression plasmids

A panel of 20 alanine-scanning mutants were created in the pN3-Gag plasmid, specifically in the MHR domain of the HTLV-1 CA_CTD_, by using the Gibson assembly method as previously described [50]. Selected mutants of interest were also introduced into the pEYFP-N3-Gag expression plasmid for subcellular localization. All mutants were confirmed by Sanger DNA sequencing analysis.

### Virus particle production

The efficiency of immature particle production was analyzed by quantifying Gag proteins in released particles through harvesting cell culture supernatants and by using immunoblot analysis with a mouse monoclonal anti-HTLV p24 antibody (Catalogue #: sc-53891; Santa Cruz Biotechnology, Dallas, TX). Briefly, the pN3-HTLV-1-Gag plasmid and the HTLV-1 Env expression plasmid were co-transfected into HEK293T/17 cells by using GenJet, ver II (SignaGen, Gaithersburg, MD) at a 10:1 ratio, respectively. After 48-h post-transfection, the viral supernatants were harvested, clarified by centrifugation (1,800 × *g* for 10 min), and filtered through 0.2 µm filters. Next, the filtered cell culture supernatants were concentrated by ultracentrifugation in a 50.2 Ti rotor (Beckman Coulter, Brea, CA) at 35,000 rpm for 90 min through an 8% Opti-prep (Sigma-Aldrich, St. Louis, MO) cushion. Particle pellets were resuspended in 1× STE buffer (100 mM NaCl, 10 mM Tris pH 8.0, and 1 mM EDTA) (G-Biosciences, St. Louis, MO). The 293T/17 cells were collected and lysed with RIPA lysis buffer and clarified via centrifugation (1800 × *g* for 10 min). The Gag proteins in supernatants were detected with a 1:2,000 dilution of anti-HTLV p24 antibody in 5% milk TBST (Tris-buffered saline plus Tween-20), while the Gag proteins in cell lysates were detected with 1:1,500 dilution of anti-HTLV p24 antibody in 5% milk TBST. Glyceraldehyde 3-phosphate dehydrogenase (GAPDH) in cell lysate was detected by using a 1:1,000 anti-GAPDH hFAB™ Rhodamine antibody (Catalogue #: 12004167; Bio-Rad, Hercules, CA) in 5% milk TBST (Tris buffer saline plus Tween-20). Membranes were washed prior to incubation with a goat anti-mouse StarBright™ Blue 700 secondary (Catalogue #: 12004158; Bio-Rad, Hercules, CA). Gag expression levels were normalized relative to GAPDH levels, and the mutant Gag expression levels were determined relative to that of WT HTLV-1 Gag.

The membranes from immunoblot analyses were imaged by using a ChemiDoc Touch system (Bio-Rad, Hercules, CA) and analyzed with ImageJ. Results were analyzed by using GraphPad Prism 6.0 (GraphPad Software Inc., San Diego, CA). Relative significance between a mutant and WT was determined by using an unpaired t-test. Immunoblot analyses were done by conducting three independent replicates.

### Cryo-EM analysis of particle morphology

The pN3-HTLV-1-Gag plasmid and the HTLV-1 Env expression plasmids were co-transfected into HEK293T/17 cells by using GenJet, ver II, at a 10:1 molar ratio as previously described [51, 52]. At 48-h post-transfection, cell culture supernatants were harvested and centrifuged at 1,800 × *g* for 10 min and followed by passing through a 0.2 µm filter. Particles were then concentrated by ultracentrifugation in a 50.2 Ti rotor at 35,000 rpm for 90 min through an 8% Opti-prep cushion. Particle pellets were resuspended in about 200 µl of STE buffer prior to ultracentrifugation through a 10% to 30% Opti-Prep step gradient at 45,000 rpm for 3 h. The particle band was removed from the gradient by puncturing the side of the thin wall ultra-clear centrifuge tube (Beckman Coulter, Brea, CA) with a hypodermic syringe needle and pelleted in STE buffer at 40,000 rpm for 1 h by using an SW55 Ti rotor. The resulting virus particle pellet was then resuspended in approximately 10 µl STE buffer and frozen at -80°C until analyzed by cyro-EM.

Cryo-EM analysis of particles was done as previously described [21, 51-53]. Briefly, particle samples were thawed on ice, and approximately 3.0 µL of purified virus particles were applied to glow-discharged Quantifoil R1.2/1.3 400-mesh holey carbon-coated copper grids. The grids were then manually blotted by a piece of filter paper and plunge-frozen in liquid ethane. The frozen grids were stored in liquid nitrogen until imaging analysis.

Cryo-EM analysis was done using a Tecnai FEI G2 F30 FEG transmission electron microscope (FEI, Hillsboro, OR) at liquid nitrogen temperature operating at 300 kV. Images were recorded at a nominal magnification of 12,000× magnification under ∼15 electrons/Å2 conditions at 5 to 10 μm under-focus by using a Gatan K2 Summit direct electron detector (Gatan Inc., Pleasanton, CA). At least 250 individual immature particles of each mutant were measured using ImageJ software. Two perpendicular diameters were measured and averaged for each particle. Particle morphologies were qualitatively characterized.

### Gag subcellular distribution analysis

Gag subcellular distribution was analyzed by quantifying the degree of Gag puncta formation in cells by using confocal laser scanning microscopy to localize Gag-eYFP as previously described [21]. Briefly, HeLa cells were cultured in six-well plates on 1.5 standard glass coverslips coated with poly-L-lysine as previously described [21, 52]. HeLa cells were transiently transfected with EYFP-tagged Gag plasmids by using GenJet, ver II (SignaGen, Gaithersburg, MD). At 24-h post-transfection, cells were stained by using DAPI (Thermo Fisher Scientific, Waltham, MA) and ActinRed 555 (Invitrogen, Waltham, CA) after fixation with 4% paraformaldehyde (Thermo Fisher Scientific, Waltham, MA). Cells were imaged by using a Zeiss LSM 700 confocal laser scanning microscope with a Plan-Apochromat 63×/1.40-numeric-aperture (NA) oil objective at 1.2X zoom (Carl Zeiss, Oberkochen, Germany). At least 5 individual cells were imaged from three independent experimental replicates, for a total of 15 cells for each Gag mutant and WT Gag. The degree of Gag protein assembly into puncta was quantified by evaluating the area associated with Gag puncta divided by the total area of Gag fluorescence, which resulted in a relative measurement of puncta formation.

### Cell-free HTLV-1 CA assembly analysis

The HTLV-1 CA (Gag amino acids 132-349) was cloned into the 6×His SUMO bacterial expression plasmid (pHYRSF53, Addgene, Watertown, MA), with mutants created by using the Gibson assembly procedure. Briefly, the CA protein was expressed in BL21 (DE3) RIP pLysS *E. coli* grown in 1 L of ZY-auto induction media in a platform shaker for 16 h at 37°C [54]. Cells were pelleted and then resuspended in 50 mL lysis buffer per 1 L culture (200 mM NaCl, 50 mM Tris pH 8) and then exposed to flash freezing to aid in cell lysis. After thawing, cells were lysed by adding 10 mg lysozyme, 0.1% Triton (v/v), and sonicated for 10 sec on/off cycles at 40% amplitude for three, 5 min treatments to reduce viscosity. The cell lysate was clarified by centrifugation 12,000 × *g* for 40 min. The cell lysate was then added to 1 mL of a Ni bead slurry (Ni Sepharose^®^ High Performance; Catalogue #: 17-5268-02; Sigma-Aldrich, St. Louis, MO) and incubated overnight, shaking at 4°C. Beads were washed with 30 mL lysis buffer, 30 mL of wash buffer (200 mM NaCl, 50 mM Tris pH 8, 50 mM imidazole) and eluted in 5 mL of elution buffer (200 mM NaCl, 50 mM Tris pH 8, 500 mM imidazole). The eluent was dialyzed against 1 L of dialysis buffer (200 mM NaCl, 50 mM Tris pH 8, 1 mM DTT) for 4 hours at 4°C, then swapped into fresh 1 L of dialysis buffer. Purified ULP1 protease was added to the eluent overnight at 4°C (pFGET19_Ulp1, Addgene, Watertown, MA). The cleaved product was loaded onto a Ni column (Cytvia HisTrap™ High Performance; Catalogue #: GHC-17-5247-01; Sigma-Aldrich, St. Louis, MO) and washed with buffer A (200 mM NaCl, 50 mM Tris pH 8, 5 mM imidazole) for 5 mL and then eluted over a 20 mL linear gradient of buffer B (200 mM NaCl, 50 mM Tris pH 8, 200 mM imidazole). The untagged HTLV-1 CA has a slight affinity for the Ni column, typically eluting approximately in 5-10% buffer B. Protein-containing fractions were run on SDS-PAGE gels to confirm purify.

Purified fractions were concentrated to approximate 5-10 mg/ml and dialyzed into assembly buffer (50 mM MES, 50 mM NaCl, pH 6.5) overnight at 4°C in Pierce™ Slide-A-Lyzer^®^ Mini Dialysis units (Thermo Fisher Scientific, Waltham, MA). The dialyzed protein was centrifuged at 16,000 × g for 5 min to remove any aggregates. Protein was either directly used or flash frozen and stored at -80°C.

For the assembly assay, the protein was diluted to 100 µM, and 100 μL was added to a 96 -well plate. Initial readings were taken at 0 min, then 150 µM IP6 (Catalogue #: P0409; TCI America, Portland, OR) was added to initiate assembly. A reading at 1 min was taken, then spectrometry readings were collected every 5 min for 1 h using a BioTek Epoch microplate spectrophotometer (Agilent Technologies, Santa Clara, CA).

After allowing assembly to proceed for 1 h, the reactions were centrifuged at 5,000 × *g* for 5 minutes to pellet the helical assemblies. The pellet was resuspended in the assembly buffer containing 150 µM IP6 and 3 µl of the resuspension was spotted onto EMS CF300-CU grids (Ted Pella Inc., Redding, CA) for 2 min. Samples were then blotted with filter paper, washed 3 times in deionized water, blotted, and stained in 0.75% (w/v) uranyl formate for 2 min. Samples were imaged using a FEI Technai Spirit Bio-Twin transmission electron microscope at 120 kV.

## Acknowledgments

This research was supported by grants from the National Institutes of Health (NIH). TEM was conducted in the Characterization Facility, College of Science & Engineering, University of Minnesota, which receives partial support from NSF through the MRSEC and NNCI program.

## References

[1] Poiesz BJ, Ruscetti FW, Gazdar AF, Bunn PA, Minna JD, Gallo RC. Detection and isolation of type C retrovirus particles from fresh and cultured lymphocytes of a patient with cutaneous T-cell lymphoma. Proc Natl Acad Sci U S A. 1980;77:7415–9.

[2] Yoshida M, Miyoshi I, Hinuma Y. Isolation and characterization of retrovirus from cell lines of human adult T-cell leukemia and its implication in the disease. Proc Natl Acad Sci U S A. 1982;79:2031–5.

[3] Kalyanaraman VS, Sarngadharan MG, Robert-Guroff M, Miyoshi I, Golde D, Gallo RC. A new subtype of human T-cell leukemia virus (HTLV-II) associated with a T-cell variant of hairy cell leukemia. Science. 1982;218:571–3.

[4] Calattini S, Chevalier SA, Duprez R, Bassot S, Froment A, Mahieux R, et al. Discovery of a new human T-cell lymphotropic virus (HTLV-3) in Central Africa. Retrovirology. 2005;2:30.

[5] Calattini S, Chevalier SA, Duprez R, Afonso P, Froment A, Gessain A, et al. Human T-cell lymphotropic virus type 3: complete nucleotide sequence and characterization of the human tax3 protein. J Virol. 2006;80:9876–88.

[6] Wolfe ND, Heneine W, Carr JK, Garcia AD, Shanmugam V, Tamoufe U, et al. Emergence of unique primate T-lymphotropic viruses among central African bushmeat hunters. Proc Natl Acad Sci U S A. 2005;102:7994–9.

[7] Uchiyama T, Yodoi J, Sagawa K, Takatsuki K, Uchino H. Adult T-cell leukemia: clinical and hematologic features of 16 cases. Blood. 1977;50:481–92.

[8] Osame M, Usuku K, Izumo S, Ijichi N, Amitani H, Igata A, et al. HTLV-I associated myelopathy, a new clinical entity. Lancet. 1986;1:1031–2.

[9] Gessain A, Barin F, Vernant JC, Gout O, Maurs L, Calender A, et al. Antibodies to human T-lymphotropic virus type-I in patients with tropical spastic paraparesis. Lancet. 1985;2:407–10.

[10] Gessain A, Cassar O. Epidemiological Aspects and World Distribution of HTLV-1 Infection. Front Microbiol. 2012;3:388.

[11] Grigsby IF, Zhang W, Johnson JL, Fogarty KH, Chen Y, Rawson JM, et al. Biophysical analysis of HTLV-1 particles reveals novel insights into particle morphology and Gag stochiometry. Retrovirology. 2010;7:75.

[12] Gillet NA, Malani N, Melamed A, Gormley N, Carter R, Bentley D, et al. The host genomic environment of the provirus determines the abundance of HTLV-1-infected T-cell clones. Blood. 2011;117:3113–22.

[13] Oliveira PD, Kachimarek AC, Bittencourt AL. Early Onset of HTLV-1 Associated Myelopathy/Tropical Spastic Paraparesis (HAM/TSP) and Adult T-cell Leukemia/Lymphoma (ATL): Systematic Search and Review. J Trop Pediatr. 2018;64:151–61.

[14] Flexner C. HIV drug development: the next 25 years. Nat Rev Drug Discov. 2007;6:959–66.

[15] Link JO, Rhee MS, Tse WC, Zheng J, Somoza JR, Rowe W, et al. Clinical targeting of HIV capsid protein with a long-acting small molecule. Nature. 2020;584:614–8.

[16] Bester SM, Wei G, Zhao H, Adu-Ampratwum D, Iqbal N, Courouble VV, et al. Structural and mechanistic bases for a potent HIV-1 capsid inhibitor. Science. 2020;370:360–4.

[17] Segal-Maurer S, DeJesus E, Stellbrink HJ, Castagna A, Richmond GJ, Sinclair GI, et al. Capsid Inhibition with Lenacapavir in Multidrug-Resistant HIV-1 Infection. N Engl J Med. 2022;386:1793–803.

[18] Tu JJ, Maksimova V, Ratner L, Panfil AR. The Past, Present, and Future of a Human T-Cell Leukemia Virus Type 1 Vaccine. Front Microbiol. 2022;13:897346.

[19] Marino-Merlo F, Balestrieri E, Matteucci C, Mastino A, Grelli S, Macchi B. Antiretroviral Therapy in HTLV-1 Infection: An Updated Overview. Pathogens. 2020;9.

[20] Maldonado JO, Cao S, Zhang W, Mansky LM. Distinct Morphology of Human T-Cell Leukemia Virus Type 1-Like Particles. Viruses. 2016;8.

[21] Martin JL, Mendonca LM, Marusinec R, Zuczek J, Angert I, Blower RJ, et al. Critical Role of the Human T-Cell Leukemia Virus Type 1 Capsid N-Terminal Domain for Gag-Gag Interactions and Virus Particle Assembly. J Virol. 2018;92:e00333–18.

[22] Mammano F, Ohagen A, Hoglund S, Gottlinger HG. Role of the major homology region of human immunodeficiency virus type 1 in virion morphogenesis. J Virol. 1994;68:4927–36.

[23] Bocanegra R, Fuertes MA, Rodriguez-Huete A, Neira JL, Mateu MG. Biophysical analysis of the MHR motif in folding and domain swapping of the HIV capsid protein C-terminal domain. Biophys J. 2015;108:338–49.

[24] Mateu MG. Conformational Stability of Dimeric and Monomeric Forms of the C-terminal Domain of Human Immunodeficiency Virus-1 Capsid Protein. Journal of Molecular Biology. 2002;318:519–31.

[25] EbbetsReed D, Scarlata S, Carter CA. The major homology region of the HIV-1 gag precursor influences membrane affinity. Biochemistry. 1996;35:14268–75.

[26] Tanaka M, Robinson BA, Chutiraka K, Geary CD, Reed JC, Lingappa JR. Mutations of Conserved Residues in the Major Homology Region Arrest Assembling HIV-1 Gag as a Membrane-Targeted Intermediate Containing Genomic RNA and Cellular Proteins. J Virol. 2016;90:1944–63.

[27] von Schwedler UK, Stray KM, Garrus JE, Sundquist WI. Functional surfaces of the human immunodeficiency virus type 1 capsid protein. J Virol. 2003;77:5439–50.

[28] Ganser-Pornillos BK, von Schwedler UK, Stray KM, Aiken C, Sundquist WI. Assembly properties of the human immunodeficiency virus type 1 CA protein. J Virol. 2004;78:2545–52.

[29] Chang YF, Wang SM, Huang KJ, Wang CT. Mutations in capsid major homology region affect assembly and membrane affinity of HIV-1 Gag. J Mol Biol. 2007;370:585–97.

[30] Koma T, Kotani O, Miyakawa K, Ryo A, Yokoyama M, Doi N, et al. Allosteric Regulation of HIV-1 Capsid Structure for Gag Assembly, Virion Production, and Viral Infectivity by a Disordered Interdomain Linker. J Virol. 2019;93.

[31] Lokhandwala PM, Nguyen TL, Bowzard JB, Craven RC. Cooperative role of the MHR and the CA dimerization helix in the maturation of the functional retrovirus capsid. Virology. 2008;376:191–8.

[32] Cairns TM, Craven RC. Viral DNA synthesis defects in assembly-competent Rous sarcoma virus CA mutants. J Virol. 2001;75:242–50.

[33] Craven RC, Leure-duPree AE, Weldon RA, Jr., Wills JW. Genetic analysis of the major homology region of the Rous sarcoma virus Gag protein. J Virol. 1995;69:4213–27.

[34] Purdy JG, Flanagan JM, Ropson IJ, Rennoll-Bankert KE, Craven RC. Critical role of conserved hydrophobic residues within the major homology region in mature retroviral capsid assembly. J Virol. 2008;82:5951–61.

[35] Strambio-de-Castillia C, Hunter E. Mutational analysis of the major homology region of Mason-Pfizer monkey virus by use of saturation mutagenesis. J Virol. 1992;66:7021–32.

[36] Andrawiss M, Takeuchi Y, Hewlett L, Collins M. Murine leukemia virus particle assembly quantitated by fluorescence microscopy: role of Gag-Gag interactions and membrane association. J Virol. 2003;77:11651–60.

[37] Rayne F, Bouamr F, Lalanne J, Mamoun RZ. The NH2-terminal domain of the human T-cell leukemia virus type 1 capsid protein is involved in particle formation. J Virol. 2001;75:5277–87.

[38] Martin JL, Cao S, Maldonado JO, Zhang W, Mansky LM. Distinct particle morphologies revealed through comparative parallel analyses of retrovirus-like particles. J Virol. 2016.

[39] Maldonado JO, Angert I, Cao S, Berk S, Zhang W, Mueller JD, et al. Perturbation of Human T-Cell Leukemia Virus Type 1 Particle Morphology by Differential Gag Co-Packaging. Viruses. 2017;9.

[40] Marie V, Gordon ML. The HIV-1 Gag Protein Displays Extensive Functional and Structural Roles in Virus Replication and Infectivity. Int J Mol Sci. 2022;23.

[41] Zurowska K, Alam A, Ganser-Pornillos BK, Pornillos O. Structural evidence that MOAP1 and PEG10 are derived from retrovirus/retrotransposon Gag proteins. Proteins. 2022;90:309–13.

[42] Gamble TR, Yoo S, Vajdos FF, von Schwedler UK, Worthylake DK, Wang H, et al. Structure of the carboxyl-terminal dimerization domain of the HIV-1 capsid protein. Science. 1997;278:849–53.

[43] Krebs AS, Mendonca LM, Zhang P. Structural Analysis of Retrovirus Assembly and Maturation. Viruses. 2021;14.

[44] Dick RA, Zadrozny KK, Xu C, Schur FKM, Lyddon TD, Ricana CL, et al. Inositol phosphates are assembly co-factors for HIV-1. Nature. 2018;560:509–12.

[45] Schur FK, Hagen WJ, Rumlova M, Ruml T, Muller B, Krausslich HG, et al. Structure of the immature HIV-1 capsid in intact virus particles at 8.8 A resolution. Nature. 2015;517:505–8.

[46] Novikova M, Zhang Y, Freed EO, Peng K. Multiple Roles of HIV-1 Capsid during the Virus Replication Cycle. Virol Sin. 2019;34:119–34.

[47] Joshi A, Garg H, Ablan S, Freed EO, Nagashima K, Manjunath N, et al. Targeting the HIV entry, assembly and release pathways for anti-HIV gene therapy. Virology. 2011;415:95–106.

[48] Dalessio PM, Craven RC, Lokhandwala PM, Ropson IJ. Lethal mutations in the major homology region and their suppressors act by modulating the dimerization of the rous sarcoma virus capsid protein C-terminal domain. Proteins. 2013;81:316–25.

[49] Khorasanizadeh S, Campos-Olivas R, Summers MF. Solution structure of the capsid protein from the human T-cell leukemia virus type-I. J Mol Biol. 1999;291:491–505.

[50] Heydenreich FM, Miljus T, Jaussi R, Benoit R, Milic D, Veprintsev DB. High-throughput mutagenesis using a two-fragment PCR approach. Sci Rep. 2017;7:6787.

[51] Martin JL, Cao S, Maldonado JO, Zhang W, Mansky LM. Distinct Particle Morphologies Revealed through Comparative Parallel Analyses of Retrovirus-Like Particles. J Virol. 2016;90:8074–84.

[52] Yang H, Talledge N, Arndt WG, Zhang W, Mansky LM. Human Immunodeficiency Virus Type 2 Capsid Protein Mutagenesis Reveals Amino Acid Residues Important for Virus Particle Assembly. J Mol Biol. 2022;434:167753.

[53] Martin JL, Mendonca LM, Angert I, Mueller JD, Zhang W, Mansky LM. Disparate Contributions of Human Retrovirus Capsid Subdomains to Gag-Gag Oligomerization, Virus Morphology, and Particle Biogenesis. J Virol. 2017;91.

[54] Studier FW. Protein production by auto-induction in high density shaking cultures. Protein Expr Purif. 2005;41:207–34.

